# A computational model of the DNA damage-induced IKK/ NF-κB pathway reveals a critical dependence on irradiation dose and PARP-1

**DOI:** 10.1101/2023.05.02.538686

**Authors:** Fabian Konrath, Michael Willenbrock, Dorothea Busse, Claus Scheidereit, Jana Wolf

## Abstract

The activation of IKK/ NF-κB by genotoxic stress is a crucial process in the DNA damage response. Due to the anti-apoptotic impact of NF-κB it can affect cell fate decisions upon DNA damage and therefore interfere with tumour therapy-induced cell death. Here, we developed a dynamical model describing IKK/ NF-κB signalling that faithfully reproduces quantitative time course data and enabled a detailed analysis of pathway regulation. The approach elucidates a pathway topology with two hubs, where the first integrates signals from two DNA damage sensors and the second forms a coherent feedforward loop. The analyses reveal a critical role of the sensor protein PARP-1 in the pathway regulation. Introducing a method for calculating the impact of individual components on pathway activity in a time-resolved manner we show how irradiation dose influences pathway activation. Our results give a mechanistic understanding relevant for the interpretation of experimental and clinical studies.

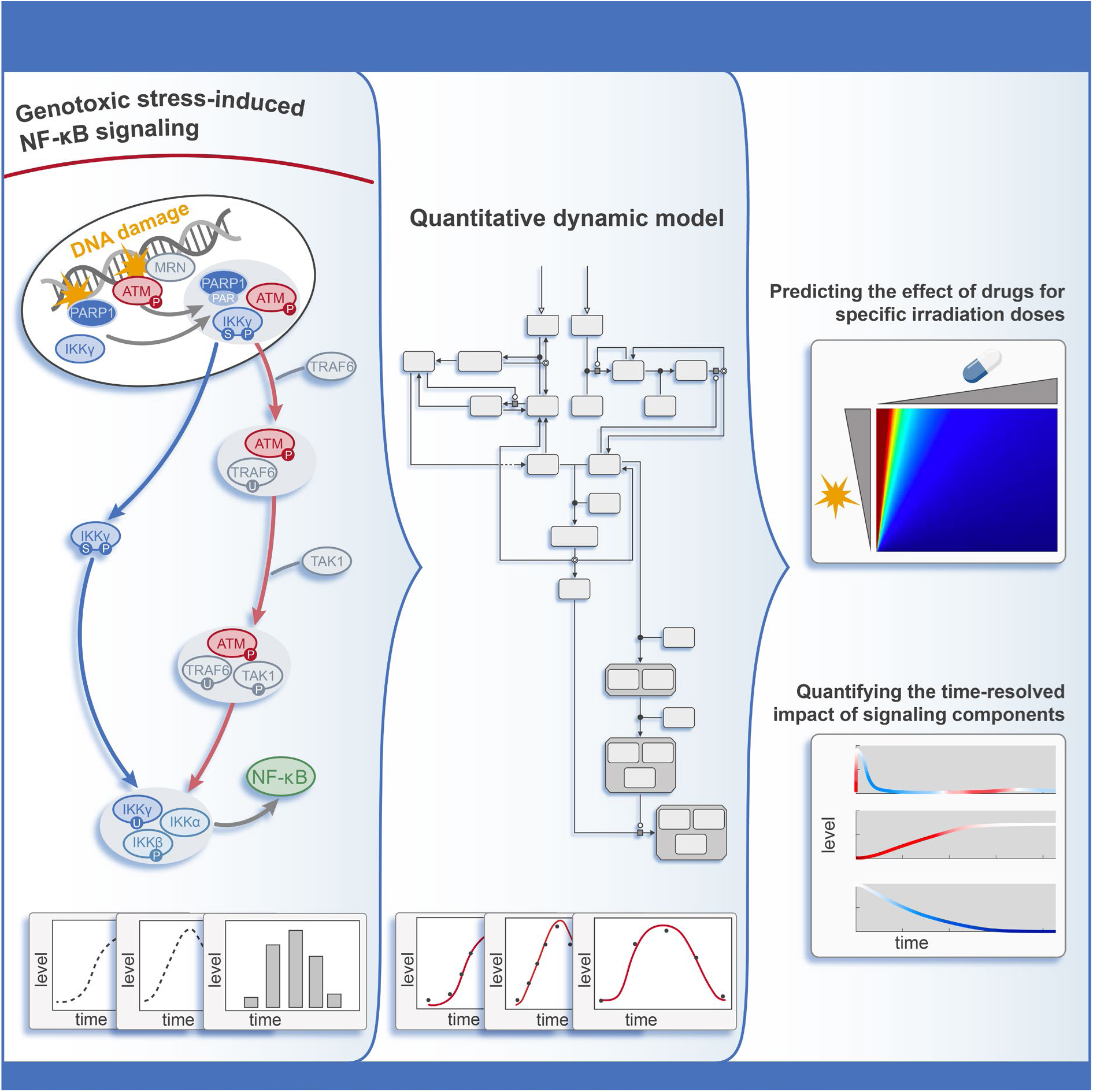

## Introduction

The members of the NF-κB transcription factor family are involved in the regulation of important cellular processes such as proliferation and survival. Over the past decades, numerous studies revealed the fundamental role of NF-κB in the regulation of inflammation, immunity as well as the cell fate decision between survival and apoptosis. Based on the stimulus and the involved components, NF-κB signalling can be divided in three pathways: The canonical, non-canonical and genotoxic stress-induced signalling cascade. While cell surface receptors such as cytokine or antigen receptors trigger canonical and non-canonical signalling, genotoxic stress signals, referred to here primarily as induced by genomic DNA double strand breaks, emanate from the nucleus.

Despite their differences, all three signalling systems culminate in the activation of NF-κB transcription factors in various crucial cellular processes, therefore underscoring the detrimental consequences of dysregulated NF-κB activity. Auto-immune diseases like psoriasis or the survival of cancer cells are examples for hyperactivated NF-κB ^1^. Hence, the biochemical interaction networks involved in the regulation of NF-κB activation were subject of numerous studies. For the canonical pathway and parts of the non-canonical pathway, computational models were developed and their systematic and detailed analyses greatly contributed to the understanding of the regulation of the NF-κB family members ^2^. In particular, multiple ODE models of the canonical pathway were developed to study the dynamical behaviour of NF-κB ^3–5^, its regulation via IκBs and A20 ^6–10^ as well as its role in the B-cell development ^11^. For the non-canonical pathway, ODE models were used to elucidate the impact of NF-κB precursors and their processing on the regulation of NF-κB signalling ^12, 13^.

For the DNA damage-induced signalling pathway, seminal experimental work revealed the signalling components involved in transferring the signal emerging from DNA lesions to the activation of NF-κB and the concomitant expression of its target genes (reviewed in ^14^). In general, a cell is permanently challenged by various kinds of external and internal stresses. For instance, UV irradiation or reactive oxygen species are genotoxic stresses that can damage DNA and thereby cause genomic instability ^15, 16^. To prevent malignant transformation, cells need to respond to such DNA lesions in an appropriate way. The DNA damage response comprises recognition of damaged DNA, signal transduction and the final cellular response ^17^. The processes and pathways that are involved in the signal transduction are dependent on the damage type. The most severe damage are DNA double strand breaks (DSBs). Homologous recombination and non-homologous end joining are two pathways that allow the repair of DSBs. However, in case of complex or irreparable damage, apoptosis or a permanent cell cycle arrest program termed senescence is induced ^18, 19^. The p65/p50 heterodimer (from here on referred to as NF-κB) is a member of the NF-κB transcription factor family and is activated during the DNA damage response ^14, 20^. Similar to the canonical pathway, the activity of NF-κB is tightly controlled by IKK, a complex consisting of kinase IKKα (IKK1), kinase IKKβ (IKK2) and the regulatory subunit IKKγ (NEMO) ^21^. Upon activation, IKKβ induces the activation of NF-κB by mediating the phosphorylation and proteasomal degradation of IκB proteins, inhibitors of NF-κB, that sequester NF-κB in the cytoplasm ^22^. Moreover, IKKβ mediates posttranslational modifications of p65 which are crucial for full activation of NF-κB ^21, 23^. For genotoxic stress-induced IKK/ NF-κB signalling, IKKγ plays a central role by transferring the signal from the nucleus to the cytoplasm ^14^.

First, the sensors poly(adenosine diphosphate (ADP)-ribose) polymerase 1 (PARP-1) and the MRN complex, consisting of MRE11, RAD50 and NBS1, detect DNA double strand breaks. The MRN complex activates the kinase ATM ^24, 25^. Activated PARP-1 and ATM form a nuclear complex with IKKγ, which leads to the posttranslational modifications of IKKγ in the nucleus ^26^. The genotoxic stress-induced signal is then transferred to the cytoplasm by shuttling of modified IKKγ to the cytoplasm and its integration into IKK complexes ^14, 27, 28^. In parallel, phosphorylated ATM translocates to the cytoplasm and mediates the formation of a TRAF6 and TAK1 containing complex which in turn facilitates the activation of the primed IKK complexes by the phosphorylation of the IKKβ subunit ^28^. The activated IKK complex phosphorylates IκBα and thereby mediates the proteasomal degradation of the NF-κB inhibitor ^22^. Freed NF-κB can translocate into the nucleus, where it induces the expression of various target genes encoding anti-apoptotic factors such as XIAP, c-FLIP, BCL-XL, and BCL2 ^29–32^ that counteract apoptosis and thereby promote cell survival ^33^.

Due to its impact on cell fate decisions, aberrant functionalities of NF-κB can result in tumour development as well as tumour progression ^34^. Activated NF-κB is observed in several types of cancer ^35, 36^ and may mediate resistance to apoptosis, a known critical event in tumorigenesis ^16^. Importantly, the choice between cell death and survival upon DNA damage is also of great importance for tumour therapy as a common therapeutic strategy is to induce DNA damage and promote cell death of tumour cells ^37, 38^. Hence, chemotherapy as well as radiotherapy trigger the activation of NF-κB ^39–41^ which in turn may interfere with therapy-induced cell death. Consequently, blocking NF-κB activity can enhance the susceptibility of tumour cells to therapy ^42, 43^.

Hence, genotoxic stress-induced IKK/ NF-κB signalling is of importance to advance the understanding of tumour development and improved therapeutic strategies. To gain mechanistic insights into oncogenic signalling pathways, quantitative computational models have been shown to be powerful tools ^44, 45^.

Here, we developed a dynamical model of the IKK/ NF-κB activation induced by DNA damage. The model, consisting of a system of coupled ordinary differential equations, is based on the biochemical network described in the experimental literature. For most of the components involved in genotoxic stress signalling, profound time-resolved data exist for different irradiation doses and cell types used in the different studies. We integrated the ample experimental observations on a quantitative level and were able to create a computational model that faithfully describes IKK/ NF-κB signalling in tumour cell lines. We then used the model for a detailed analysis of the regulation of IKK/ NF-κB signalling triggered by DNA damage. Introducing a new method that allows to assess the impact of individual signalling components on a process in a temporally resolved and systematic manner we could show that the regulation of the signalling is a highly dynamic process that depends on the irradiation dose. The analysis allowed the identification of potential drug targets with a major impact on inhibiting the activation of the IKK complex for different irradiation doses.

## Results

### A mathematical model describing the activation of IKK/ NF-κB signalling by genotoxic stress

We developed an ordinary differential equation-based model based on literature derived mechanisms and experiment-informed refinements (for details of the model development and incorporated experiments see Text S1). The model describes the initial recruitment of the DNA damage sensors PARP-1 and the MRN complex to DNA lesions, the integration of the signal, its transmission from the nucleus to the cytoplasm via IKKγ and ATM as well as the final activation of the IKK complex (see Fig 1A for a model scheme and Text S1 for model details). In presence of DSBs, the recruited MRN complex mediates the recruitment and activation of the kinase ATM ^46, 47^. PARP-1 undergoes automodification by attaching poly(ADP-ribose) (PAR) molecules to itself and thereby forms a platform for the activated ATM, the SUMO-1 ligase PIASy and IKKγ ^26^. The close proximity of the components facilitates the phosphorylation and sumoylation of IKKγ ^26, 48, 49^. The nuclear complex, termed signalosome, disintegrates due to dePARylation of PARP-1 which is mediated by the PAR glycohydrolase ^50^. The complex disassembly frees the posttranslationally modified IKKγ and enables its translocation to the cytoplasm ^14, 26, 27^ where it integrates into an IKK complex ^26^.

**Fig 1.**
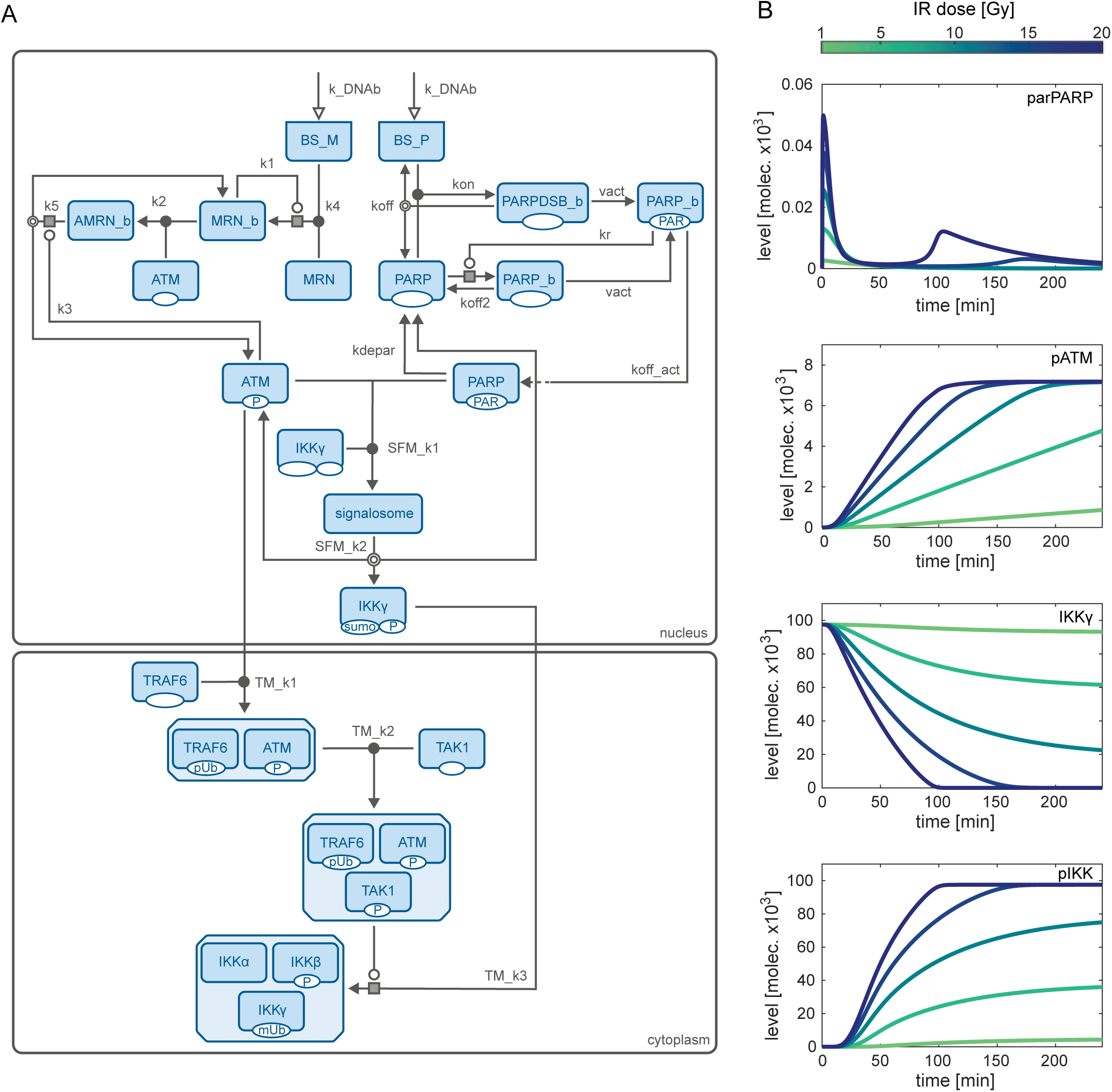
Model scheme and simulations of selected pathway components. (A) The model describes the following nuclear processes: PARylation of PARP-1 upon its recruitment to the binding site BS_P, recruitment of MRN to binding site BS_M, activation of ATM and formation of the signalosome consisting of PARylated PARP (parPARP), phosphorylated ATM (pATM) and IKKγ (IKKγ). Posttranslationally modified IKKγ (spIKKγ) and pATM translocate to the cytoplasm. There, the formation of the ATM-containing TRAF6 complex (AT) and the ATM-TAK1-TRAF6 complex (ATT) as well as the activation of the IKK complex (pIKK) are taken into account. PARP-1 can either bind directly to the lesion (PARPDSB_b) or it is recruited by PARylated and DNA bound PARP-1 molecules (PAR-PARP_b) to the proximity of the lesion (PARP_b). The MRN complex is recruited to the lesion (MRN_b) and in turn recruits ATM (AMRN_b). Based on the SBGN convention ^76^, posttranslational modifications are indicated below the boxes that represent network components. P: phosphorylation, PAR: PARylation, sumo: sumoylation, pUb: poly-ubiquitination, mUb: mono-ubiquitination. White arrow heads represent inputs, white circles pointing to a grey box represent catalytic processes in which a component modulates the process with the grey box, double line circles show dissociation processes and black circles represent association processes. For a detailed description of the model processes, see Text S1, Section II.A. (B) Simulation results for parPARP, pATM, IKKγ and pIKK for an irradiation dose of 1, 5, 10, 15 and 20 Gy.

Simultaneously, activated ATM translocates to the cytoplasm where it induces the formation of a complex containing TRAF6 and TAK1, which in turn mediates the activation of the primed IKK complexes ^28^. We trained the model on various data sets capturing for multiple irradiation doses detailed time course measurements of critical pathway components, such as PARP-1, ATM and IKKγ (Text S1, Section I). Using those data sets with around 10000 data points enabled us to create a model that faithfully reproduces the dynamics of genotoxic stress-induced IKK/ NF-κB signalling (for a comparison of experimental data and simulated dynamics see Fig S1). In this way, the model integrates the information of these different data sets and links them based on the underlying mechanistic network of biochemical interactions. The trained model can be used to simulate and predict the time courses of all implemented network components for various irradiation doses. Fig 1B shows the dynamics of selected pathway components: posttranslationally modified PARP-1 (parPARP), phosphorylated ATM (pATM), unmodified IKKγ (IKKγ) in the nucleus, and activated cytoplasmic IKK complex (pIKK) consisting of IKKα, phosphorylated IKKβ und posttranslationally modified IKKγ. The simulations reveal different activation dynamics of the two DNA damage sensors. While PARP-1 is fast and transiently modified (parPARP) with a peak amplitude depending on the irradiation dose, the levels of pATM increase until a constant level is reached. The slope of the increase thereby strongly depends on the irradiation dose with a faster increase for higher irradiations. Unmodified IKKγ (IKKγ) is declining over time, the levels of activated IKK complex (pIKK) increase over time. For both components, the slope and activated steady state levels change with irradiation levels. A full activation of IKK complex is reached for around 13 Gy and higher (Fig 2A).

**Fig 2.**
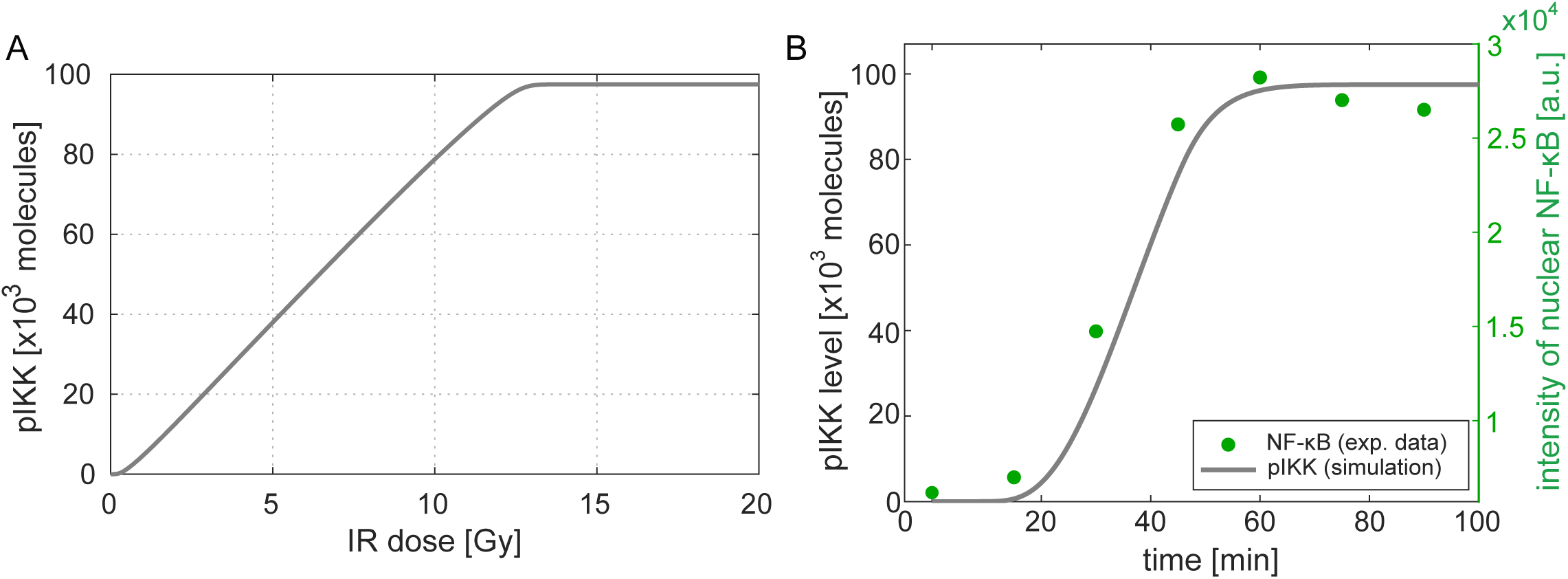
Input – output behaviour of the model and comparison of simulated IKK complex activation and experimentally quantified NF-κB activation. (A) For different irradiation doses (input), the corresponding stimulated steady state level of activated IKK complex (pIKK, output) is simulated. (B) The green dots depict quantified EMSA data of nuclear NF-κB detected in HepG2 cells after 40 Gy irradiation. The grey line represents the simulated time course of IKK complex activation. The experimental EMSA data was generated using standard procedures as described in Stilmann et al., ^26^.

To validate the model, we compared the simulated time course of activated IKK complex and measured levels of nuclear NF-κB upon 40 Gy γ-irradiation, a data set that was not used for parameter estimation. We find a good agreement between simulation and this experimental data set (Fig 2B), which demonstrates that i) the level of activated IKK complex is a good approximation for NF-κB activity and ii) the model is able to successfully reflect the dynamics of NF-κB activity upon γ-irradiation. We therefore define for the following analyses activated IKK complex as a readout (see Text S1, Section II.F for details).

### Model predicts PARP-1 inhibition as most effective target to reduce IKK complex activity

Due to the anti-apoptotic activity of NF-κB, it can be beneficial to inhibit its activation upon tumour therapy and thereby render tumour cells sensitive to therapy-induced apoptosis ^36^. Thus, we sought to identify processes that would allow to effectively interfere with the activation of IKK/ NF-κB. We therefore performed a sensitivity analysis on all model parameters for a broad range of irradiation doses ranging from 1 to 100 Gy. The parameters were decreased by 90% with respect to their nominal value and the change in the stimulated steady state of activated IKK complex (pIKK) was quantified (Fig 3A).

**Fig 3.**
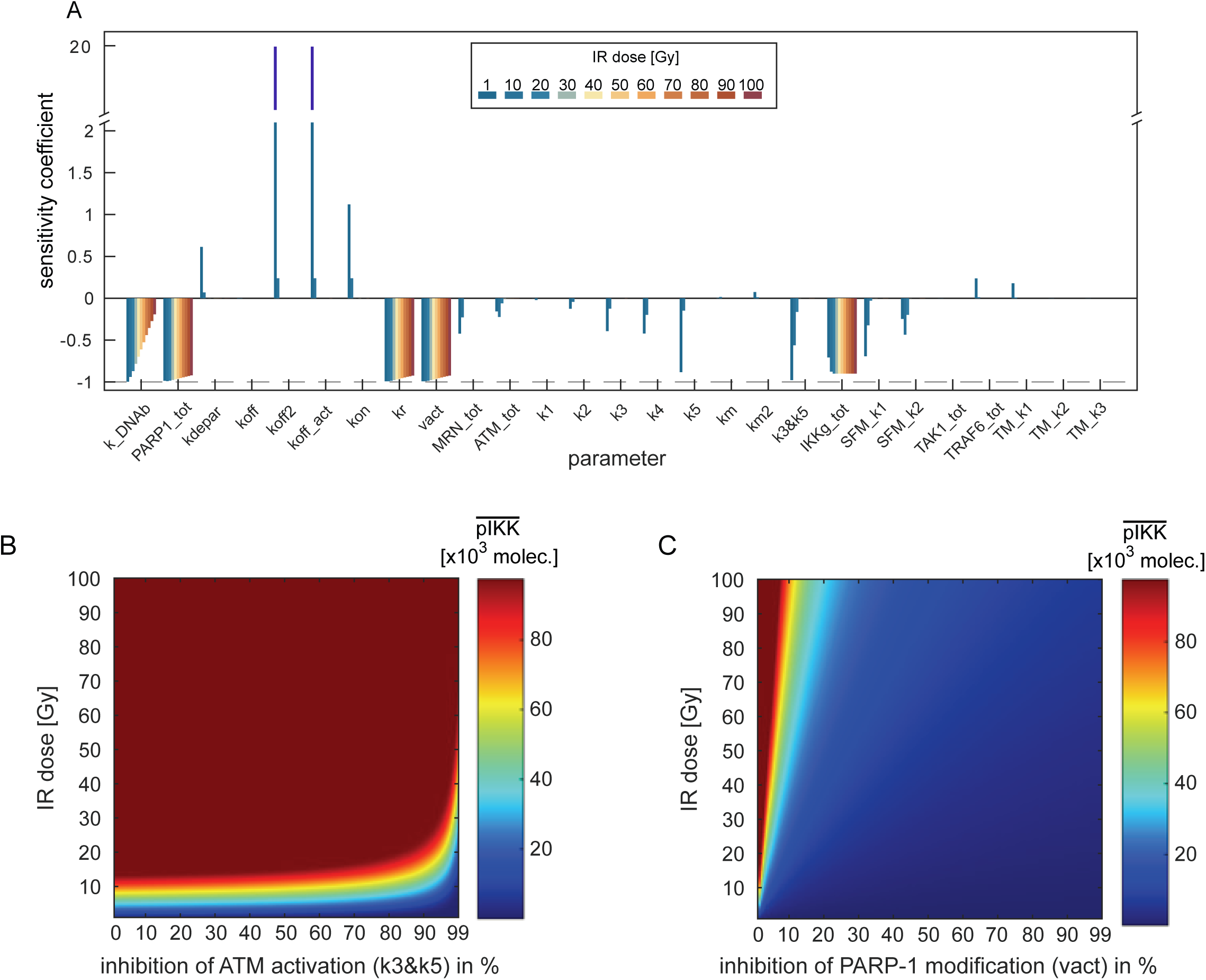
Impact of parameter perturbations on the level of activated IKK complex for various irradiation doses. (A) The values of the specified parameters were reduced by 90% for various irradiation doses. The effect of a parameter perturbation on the stimulated steady state level of the activated IKK complex is given by the sensitivity coefficient. Minus one (dashed line) indicates the maximal inhibition of IKK complex activation resulting in an activated IKK complex level of zero after parameter perturbation. Parameters with the suffix *tot* represent the total amount of a conserved moiety. To quantify the effect of ATM inhibition, an additional sensitivity coefficient was calculated for the two parameters representing ATM activation (k3 and k5, denoted by k3&k5). Both parameters were simultaneously reduced by 90%. (B) and (C) ATM activation and PARylation of PARP-1, respectively, are inhibited for various efficiencies ranging from 0% inhibition (unperturbed) to 99% inhibition. The colour in the plots indicate the amount of activated IKK complex (pIKK) molecules in the stimulated steady state. The inhibition of ATM activation is simulated by a simultaneous perturbation of the two parameters k3 and k5, the inhibition of PARylation of PARP-1 by perturbation of the parameter vact.

The parameters with the strongest negative impact on IKK activity for all tested irradiation doses are the total amount of PARP-1 (PARP1_tot) and the rate constants of PARP-1 recruitment (kr) and PARP-1 auto-modification (vact). The total amount of IKKγ (IKKg_tot) also has a negative impact on IKK complex activity for all tested irradiation doses but to a lower extent than the parameters affecting PARP-1 modification. The dissociation rate parameters koff2 and koff_act (characterizing the dissociation of PARP-1 from DNA) have a strong positive effect on the level of activated IKK complex as a reduction of those parameters causes an increase in the amount of modified PARP-1 and a stronger IKK complex activation. Interestingly, for the parameters koff2, koff_act, but also others, the sensitivity coefficients strongly depend on the irradiation dose. This is also true for an inhibition of ATM activation, represented in Fig 3A by parameter label k3&k5, which strongly reduces the level of activated IKK complex for 1 Gy, but has only a minor effect for higher irradiation doses (Fig 3A). To analyse the dependency of the sensitivities on the applied irradiation dose in more detail, we focused on the effect of two processes, namely activation of ATM (k3&k5) and PARylation of PARP-1 (vact), which are targets of clinically relevant inhibitors ^51, 52^. We simulated the level of activated IKK complex in the stimulated steady state for different inhibition efficiencies of ATM activation and PARP-1 modification as well as for various irradiation doses (Fig 3B and Fig 3C). With unperturbed ATM activity (inhibition efficiency: 0%), the amount of activated IKK complex increases with increasing irradiation doses until the maximal level of activated IKK is reached (Fig 3B). Inhibition of ATM activity by 90% reduces the level of activated IKK, however, this is only valid for irradiation doses below 25 Gy. For higher irradiation doses, a 90% inhibition of ATM activity is not sufficient to decrease the level of activated IKK complex. In order to reduce the level of activated IKK complex in the range of high irradiation doses, the inhibition efficiency of ATM activity needs to be increased. For example, for an irradiation dose of around 80 Gy even a 99% inhibition of ATM activity has only a negligible effect on the IKK complex activity. In contrast, an inhibition of PARP-1 auto-modification has for all irradiation doses a much stronger effect on the level of activated IKK complex (Fig 3C). Even for an irradiation dose of 100 Gy, inhibition of PARP-1 auto-modification by 50% is sufficient to strongly reduce the amount of activated IKK complex. Consequently, these results reveal that activation of IKK complex and thus the activity of NF-κB is more sensitive to PARP-1 inhibition than to ATM inhibition.

### Elucidating the time-resolved impact of pathway components uncovers an irradiation dose-dependent shift in critical components

To mechanistically understand the results of the sensitivity and inhibitor analysis, we investigated the regulation of the pathway activation and its dependence on the irradiation dose in more detail. The derived pathway model for IKK/ NF-κB activation exhibits two hubs at which two signalling branches converge, respectively (see Fig 1). The first hub is the formation of the signalosome which connects the branches of PARP-1 modification and ATM phosphorylation. At the second hub, the IKK complex is activated by posttranslationally modified IKKγ (spIKKγ) and pATM-mediated complex formation in the cytoplasm.

We here focus on the first hub, the formation of the signalosome. Within the signalosome, IKKγ becomes posttranslationally modified, which makes that hub a critical step in the transduction of the signals. In order to understand the regulation of the signalosome formation in a time-resolved manner, we analysed the impact of each of the three components, unmodified IKKγ, parPARP and pATM, on the flux of signalosome formation (eq. 1). We therefore developed a new method that allows to determine the time-resolved impact of components on a given flux by calculating the normalized derivative of a flux (eq. 2). This way, the change in the flux is composed of the normalized change of each component and allows to assign the impact of individual changes in the components to the change in the flux at a given time point. To quantify this impact, we defined the time-resolved impact fraction *ω* for each component (eq. 3, 4 and 5).

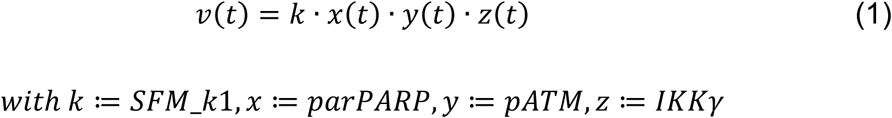

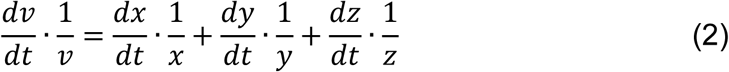

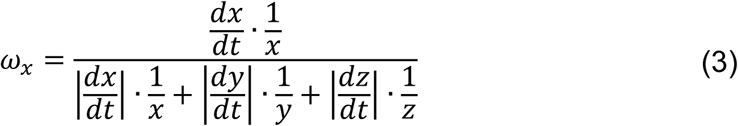

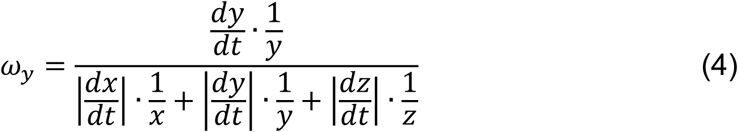

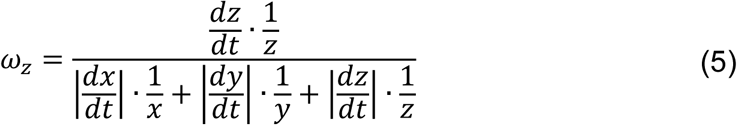

Note that the absolute values of the impact fractions of the three components sum up to one, thereby allowing to compare their impact on the derivative of flux *v*.

We first analysed the signalosome formation for an irradiation dose of 15 Gy which leads to a full activation of the IKK complex (Fig 2A). Fig 4A shows the signalosome formation flux over time, Fig 4B the dynamics of PARylated PARP-1, phosphorylated ATM and IKKγ. The impact of each component at a given time point is calculated according to eqs. (3)-(5) and the value for fraction *ω* is colour coded for each component at the corresponding time point and overlayed with its time course (Fig 4B). Comparing the trajectories in Fig 4B with the flux of signalosome formation (Fig 4A) reveals that the increase in the flux of signalosome formation within the first minutes is caused by both, the increase in the level of parPARP and pATM, which is visualized by the red colour of the corresponding trajectories. After the initial increase of parPARP, it rapidly decreases and negatively effects the flux of signalosome formation, indicated by the blue colour. However, the flux of signalosome formation further increases during the first 20 minutes since the levels of pATM positively affect the flux while the decrease in IKKγ levels do not affect it given by the white colour of the trajectory of IKKγ. Thus, after the initial increase of parPARP, pATM is the only component positively affecting the flux during this time period and is therefore solely responsible for the flux to further increase. After around 20 minutes, the flux of signalosome formation starts to decrease and eventually becomes zero (at around 190 minutes). Within this time frame, parPARP has initially a negative impact on the flux and based on the darker blue colour compared to that of IKKγ, it drives in the beginning the decrease of the flux. However, at around 100 minutes, the levels of parPARP show another, but minor increase resulting in a positive impact on the flux that counteracts, together with the positive impact of pATM, the negative impact of IKKγ. Between 20 and 190 minutes, the negative influence of IKKγ becomes stronger and ultimately IKKγ becomes the critical component determining the decrease of the flux which finally leads to the abrogation of signalosome formation.

**Fig 4.**
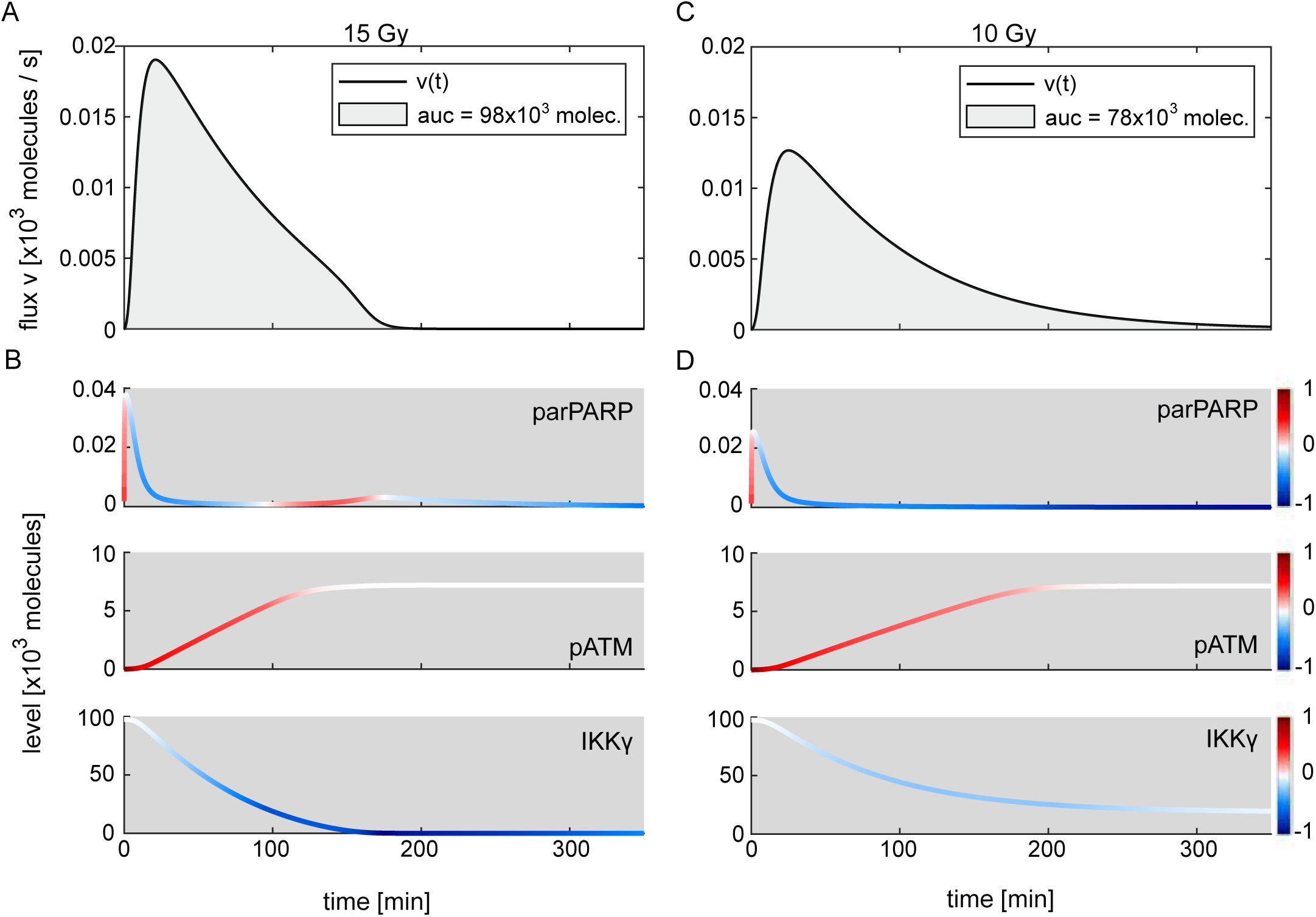
Time-resolved influence of parPARP, pATM and IKKγ on the signalosome formation flux. (A) The black line shows the simulated time course of the flux of signalosome formation for 15 Gy irradiation. The area under the curve (auc) of the flux (filled in grey) corresponds to the total number of modified IKKγ molecules. (B) The trajectories of PARylated PARP-1 (parPARP), phosphorylated ATM (pATM) and IKKγ are coloured based on the impact fraction *ω* of the respective component at a given time point. The value of an impact fraction can range from plus one and minus one. While red coloured parts of a trajectory represent a positive value and a positive impact on the flux, blue represents a negative value and a negative impact, white shows a negligible impact. (C) and (D) Simulated time course of signalosome formation upon 10 Gy and the corresponding trajectories of parPARP, pATM and IKKγ.

In order to analyse the regulation of the pIKK complex level for irradiation doses that do not lead to a maximal activation of the system, we computed the time-resolved impact fractions for the signalosome formation for 10 Gy (Fig 4C and 4D). For this lower irradiation dose, the flux of signalosome formation is lower and while the flux increase has a similar timing, the decrease of the flux takes much longer. Comparing the dynamics of parPARP, pATM and IKKγ reveals slight changes in concentrations and timings. Importantly, the impact fractions of parPARP and IKKγ show strong changes between the two irradiation doses. In contrast to the 15 Gy irradiation, the level of IKKγ does not affect signalosome formation which is indicated by a time-resolved impact fraction close to zero throughout the whole time frame of 350 minutes (IKKγ, Fig 4D). Similar to the 15 Gy irradiation scenario, the initial increase in the parPARP level positively contributes to the increase in the signalosome formation together with the increase in the pATM level. While parPARP decreases, pATM further increases and drives the increase in the flux. However, the impact of decreasing parPARP levels on the flux rises and becomes close to minus one which finally leads to the abrogation of signalosome formation.

Overall, the analyses demonstrate that the initial signalosome formation is positively impacted by parPARP and pATM, enabling an integration of signals via the two branches of DNA damage sensor signalling. The abrogation of signalosome formation is controlled by parPARP and IKKγ. Here the main control differs for different irradiation doses. While for 15 Gy and above (leading to full IKK complex activation) IKKγ essentially prevents further signalosome formation, for 10 Gy parPARP has the main control on signalosome formation.

Elucidating the control properties of signalosome formation is important as it determines the amount of IKKγ that can be posttranslationally modified during the signalling response. From the model structure one can derive that the amount of modified IKKγ (spIKKγ) determines the level of activated IKK complex in the stimulated steady state (Text S1, Section II.E). This can be demonstrated by integrating the flux of signalosome formation, which corresponds to the total amount of modified IKKγ in the nucleus over time (grey areas under the curves, Fig 4A and 4C). The calculated molecule numbers for 15 Gy and 10 Gy equal the level of activated IKK complex in steady state for the corresponding irradiation doses (Fig 2A). Consequently, the irradiation dose affects the regulation of IKK activity by causing a switch in critical components that restrict signalosome formation.

### ATM inhibition modulates the availability of PARylated PARP-1 for signalosome formation

The results of the sensitivity and inhibitor analysis (Fig 3) revealed that for higher irradiation doses the level of activated IKK complex in the stimulated steady state is more sensitive to inhibition of PARP-1 related processes of PARylation (e.g. parameter vact) than ATM inhibition (parameter k3&k5). To understand the weaker effect of ATM inhibition on the level of activated IKK complex, we evaluated the effect of this inhibition on the time-resolved impact fractions of parPARP, pATM and IKKγ during signalosome formation.

Figure 5A and 5B show the dynamics of the signalosome formation flux and the trajectories of parPARP, pATM and unmodified IKKγ, respectively, for an irradiation dose of 15 Gy and a 90% inhibition of ATM activation. As expected from the inhibitor analysis shown in Fig 3B, a 90% ATM inhibition at an irradiation dose of 15 Gy causes a reduction of the level of activated IKK complex in the cytoplasm. This reduced amount of activated IKK complex is reflected by a smaller area under the curve of the flux of signalosome formation (compare Fig 4A, 98,000 species, and Fig 5A, 58,000 species).

**Fig 5.**
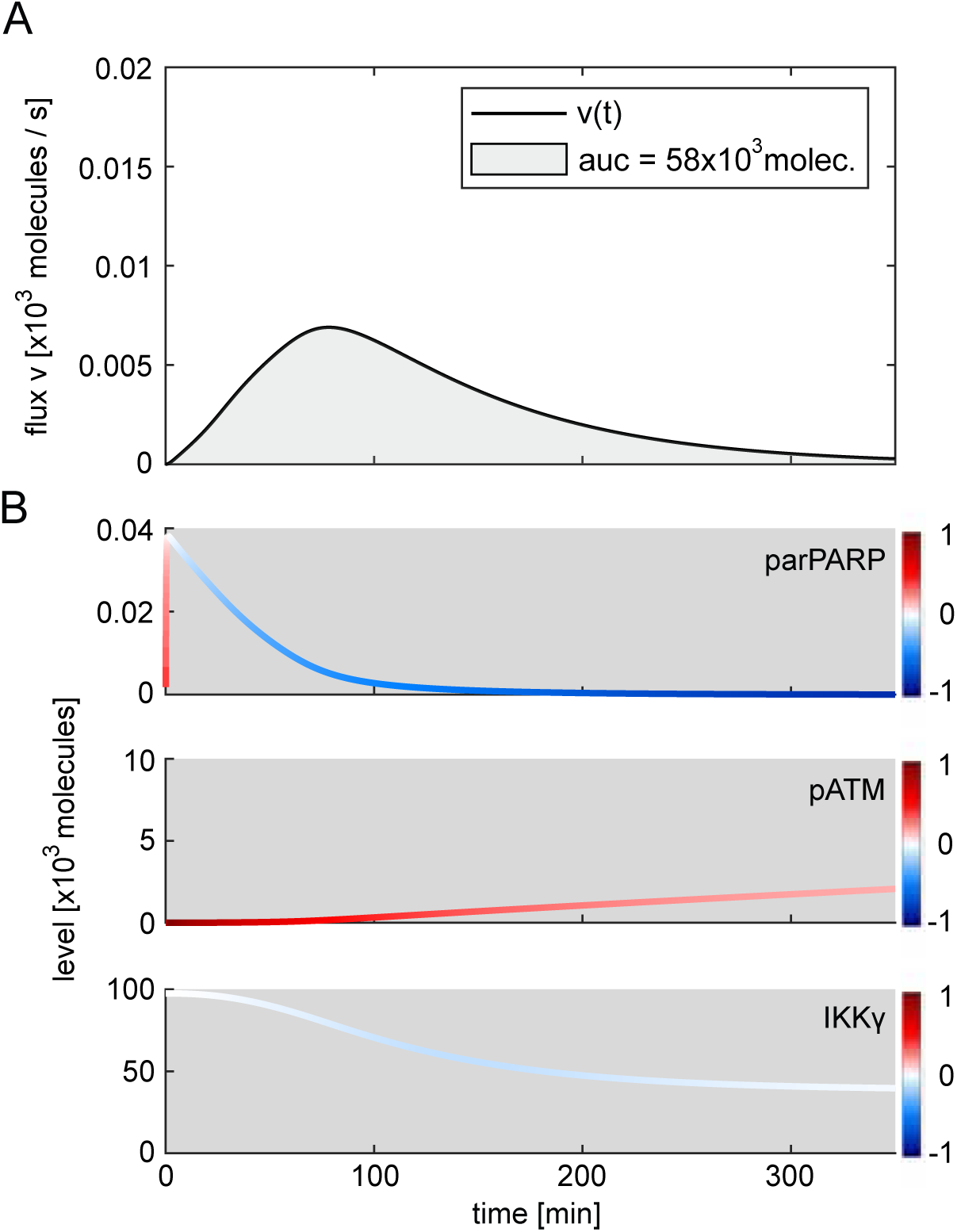
Change in the time-resolved influence of parPARP, pATM and IKKγ at 15 Gy and ATM inhibition. (A) Simulated time course of the flux of signalosome formation and area under the curve (auc) upon 15 Gy and inhibition of ATM activation by 90%. The inhibition is simulated by a simultaneous decrease in parameters k3 and k5 by 90%. (B) Trajectories of parPARP, pATM and IKKγ are coloured based on the corresponding time-resolved impact fraction ω.

The time course in Fig 5B demonstrates that an inhibition of ATM activity results in strongly reduced pATM levels during the initial increase of the flux of signalosome formation. Consequently, the increase in the flux is less steep, the maximal value is reduced and the time frame of signalosome formation is prolonged (compare Fig 5A and 4A). Strikingly, with ATM inhibition, after 80 minutes, the reduction in the level of PARylated PARP-1 becomes critical and drives the decrease of the flux of signalosome formation until the flux is abrogated (Fig 5). The decrease in the level of unmodified IKKγ also contributes to the decrease in the flux (light blue colour of the IKKγ trajectory in Fig 5B), but to a much smaller extent than the change in the parPARP level (dark blue colour of the parPARP trajectory in Fig 5B). The level of phosphorylated ATM increases during signalosome formation and has a positive impact on the flux (red colour of the pATM trajectory). However, the positive influence is counteracted by the strong negative impact of decreasing parPARP levels. Hence, inhibition of ATM activation reduces the amount of available parPARP for signalosome formation which establishes a regulatory regime, similar to the signalosome formation for an irradiation dose of 10 Gy (Fig 4D) where parPARP has the main limiting role in signalosome formation.

Based on these results, one can understand the different efficacies of ATM and PARP-1 inhibition with respect to reducing the level of activated IKK complex as it was shown in Fig 3. As can be seen in Fig 3A and B, ATM inhibition controls the level of activated IKK complex only up to a certain irradiation dose. This effect is mediated by a reduction of the availability of parPARP for signalosome formation (Fig 5). However, for higher irradiation doses, the amount of parPARP is increased (Fig 1B and Fig S2, grey line) and thereby counteracts the effect of ATM inhibition. In contrast, inhibition of PARP-1 PARylation strongly reduces the amount of parPARP for all indicated irradiation doses (Fig S2, orange line).

In summary, inhibition of ATM activation can reduce the level of activated IKK complex in the stimulated steady state by limiting the availability of PARylated PARP-1 and thereby reducing the flux of signalosome formation and consequently the amount of modified IKKγ. As increasing irradiation doses lead to higher levels of PARylated PARP-1, ATM inhibition is only effective in a certain range of irradiation doses.

### Parallel signalling branches from nucleus to cytoplasm shape the dynamics of IKK complex activation

We so far analysed the signal integration in the first hub of the pathway that is the signalosome formation in the nucleus. We now focus on the second hub, the activation of the IKK complex in the cytoplasm. Evaluating the time-resolved regulation of the second hub is of interest as it sheds light on the initiation of IKK complex activity and therefore the onset of NF-κB activity upon generation of DSBs. To assess the regulation of initial IKK complex activation, we calculated the time-resolved impact fractions of spIKKγ and the cytoplasmic TRAF6 complex (ATT) for the flux of IKK complex activation (eq. 6). We chose these two components as they have a direct impact on the activation of the IKK complex. Based on the normalized derivative of the flux (eq. 7) we calculated the impact fractions for spIKKγ, *ω_x_* (eq.8), and ATT, *ω_y_* (eq. 9).

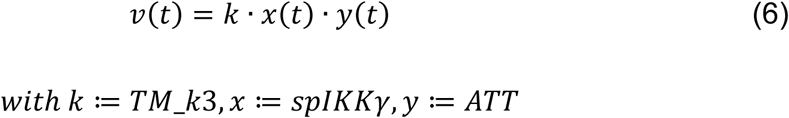

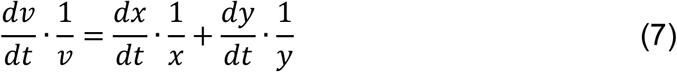

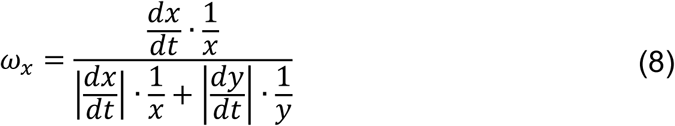

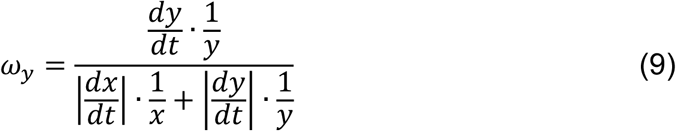

Fig 6A shows the level of activated IKK complex (pIKK, solid line) as well as the flux of IKK complex activation (dashed line) upon 15 Gy irradiation. After around 10 minutes, the level of pIKK starts to increase until it reaches the stimulated steady state level after around 200 minutes (Fig 6A, solid line). Fig 6B depicts the dynamics of spIKKγ and ATT as well as their colour coded impact fractions. The impact fractions allow to identify the critical component driving the changes in the flux of IKK complex activation. We defined a threshold of 50% for the absolute value of an impact fraction to determine which of the two components has the main impact on changes in the flux of IKK complex activation. We coloured the time frames in Fig 6A as areas under the flux accordingly. Notably, the initial increase of the flux (Fig 6A, dashed line) is positively impacted by ATT and in parts by spIKKγ. ATT is the critical component during this flux increase due to a higher impact fraction compared to spIKKγ (Fig 6A, blue area). After the critical component switches from ATT to spIKKγ which is indicated by the grey area in Fig 6A, the flux does not increase further, but starts to decrease. Since the decreasing levels of spIKKγ have a negative impact on the flux (blue coloured trajectory of spIKKγ), the dominating impact of spIKKγ counteracts the positive impact of ATT and thereby causes a decrease in the flux of IKK activation. Consequently, the level of activated IKK complex (Fig 6A, solid line) shows a reduced increase and reaches a steady state when the flux of IKK activation is zero. Hence, the initial activation of IKK complex is mediated by both components, ATT and spIKKγ, but with a main positive control of the ATM-TRAF6-TAK1-complex (ATT). The modified form of IKKγ (spIKKγ) exerts a stronger control on the level of pIKK at later time points, preventing a further activation and allowing to reach a pIKK steady state.

**Fig 6.**
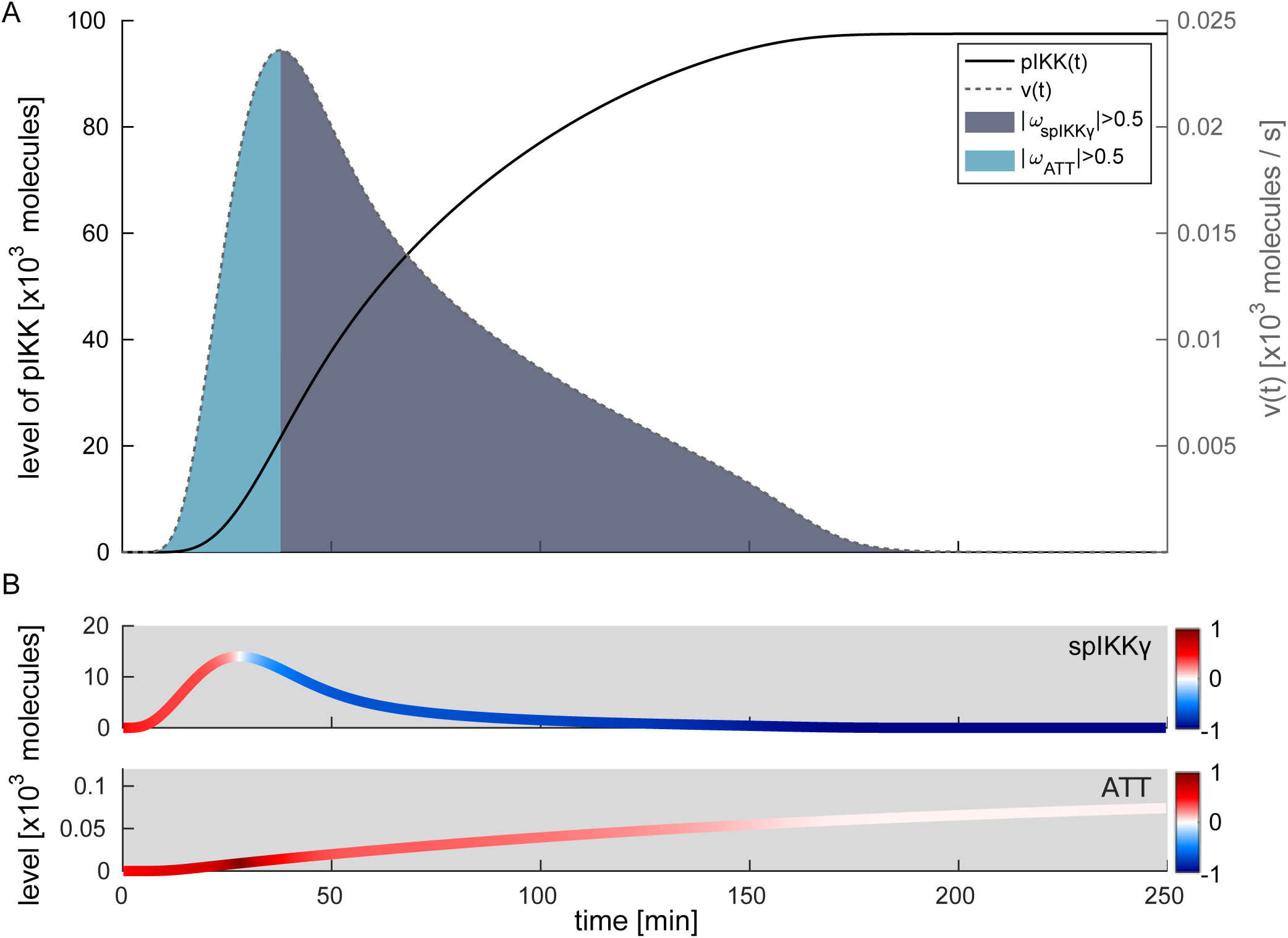
Time-resolved influence of sumoylated and phosphorylated IKKγ and the ATM-TRAF6-TAK1 complex on IKK complex activation upon 15 Gy. (A) The black solid line represents the simulated level of activated IKK complex (pIKK) over time. The flux of IKK complex activation is shown as grey dashed line. The area under the curve has a colour-code according to the component exerting the dominant impact on IKK complex activation at a given time point. The grey colour corresponds to a predominant impact of sumoylated and phosphorylated IKK (spIKKγ) and blue to a predominant impact of the ATM-TRAF6-TAK1 (ATT) complex. (B) Level of spIKKγ and ATT over time. The trajectories are coloured based on the impact fraction of the respective component at a certain time point. A red colour represents a positive impact on the flux of IKK complex activation and blue represents a negative impact.

## Discussion

Genotoxic stress-induced IKK/ NF-κB signalling plays an important role in cell fate decisions upon DNA damage and can therefore contribute to tumour development or interfere with DNA damaging tumour therapies. To gain a systematic understanding of regulatory mechanisms within the genotoxic stress-induced IKK/ NF-κB signalling network, we integrated and linked multiple data sets capturing different parts of the network from four studies ^26, 28, 53, 54^ by fitting our model to these experimental observations. This way, we developed a quantitative dynamic model describing the activation of IKK/ NF-κB by genotoxic stress. Noteworthy, additional studies exist that were not used for model development but show for parts of the activation process the involvement of additional or alternative components or interactions. For example, our model implementation of IKKγ translocation is based on the study by Hinz et al. (2010) who showed that post-translationally modified IKKγ and phosphorylated ATM translocate independently to the cytoplasm ^28^. In contrast, Jin et al., ^55^ and Wu et al., ^48^ concluded that they translocate together to the cytoplasm. Moreover, Hinz et al., ^28^ showed that auto-ubiquitination of TRAF6 leads to the formation of a complex containing IKK complexes and TAK1, while Wu et al. ^56^ and Yang et al. ^57^ reported that proteins such as RIPK1 facilitate the interaction between TAK1 and IKK complexes. Furthermore, the linear ubiquitination complex LUBAC has been implicated in the activation of TAK1 ^58^.

Despite the differences and additions, all studies report consistently that i) IKKγ is sumoylated and subsequently ubiquitinated and ii) TAK1 is activated by ATM and forms a poly-ubiquitin-based platform which promotes the activation of IKKβ and thereby NF-κB (for a detailed review, see ^14^). Of note, with our recent genome-wide screen for components of DNA damage-induced NF-κB activation, we identified scores of regulators and re-confirmed the components used in the model, including a crucial role for PARylation ^59^. Our model can therefore be seen as a core model of genotoxic stress-induced IKK/ NF-κB activation.

The computational analyses of the model enabled us to study the sensitivity of the pathway in detail. This revealed PARP-1 PARylation inhibition as the most sensitive targeting strategy to diminish the level of activated IKK complex for all considered irradiation doses. For other targets, such as phosphorylation of ATM, the inhibitory effect on IKK complex activity strongly depends on the irradiation dose. Hence, the irradiation dose and therefore the number of DNA lesions determine the sensitivity of the IKK/ NF-κB signalling pathway. Our results strongly support the notion that considering the irradiation dose is important to understand the regulation of IKK/ NF-κB signalling by DNA double strand breaks ^14, 60^. This is crucial for the evaluation and comparison of experimental and clinical studies, as the applied irradiation doses can vary greatly between them.

To assess the impact of irradiation doses on the regulation of IKK complex activity in a temporal and mechanistic manner, we developed a method that allows to quantify the impact of individual pathway components on a given flux. In general, this method can be applied to any ordinary differential equation-based model. Here, the method enabled us to investigate the impact of three pathway components, PARP-1, ATM and IKKγ, on the flux of signalosome formation and concomitantly the amount of activated IKK complex in the stimulated steady state. The results uncovered the availability of PARylated PARP-1 as a critical factor for signalosome formation. Since the level of phosphorylated ATM contributes to the flux of signalosome formation, it affects the amount of PARylated PARP-1 incorporated into the signalosome. Hence, our results indicate that the balance between phosphorylated ATM and PARylated PARP-1 is crucial for the regulation of IKK complex activity by genotoxic stress. It is therefore important to validate the quantities estimated for the model by parameter estimation (Table S2 in Text S1) with independent experimental measurements. In the study by Bakkenist and Kastan, it was shown that irradiating cells with 0.5 Gy is already sufficient to induce full activation of ATM ^47^. This is in line with our model, in which ATM is fully activated for all considered irradiation doses starting from 1 Gy. For the total amount of ATM, a value of 7·10^3^ molecules was estimated by model fitting which is in the same range as published values of quantified ATM molecules per cell ^61, 62^. The level of total PARP-1 was set in the model to a fixed value of 1.9·10^5^ molecules based on the study of Schwanhäusser et al. ^61^, which coincides with the PARP-1 molecules per cell quantified by Beck et al. ^62^. Besides ATM and PARP-1, IKKγ is also part of the signalosome and was identified in our study as the component that limits the amount of activated IKK complex at high doses of irradiation (Fig 4A and 4B). In experiments, active NF-κB, which is approximated in our model by activated IKK (Fig 2B), can be detected upon 1 Gy irradiation and was shown to saturate in two breast cancer cell lines at 20 Gy and 50 Gy, respectively ^63^. Moreover, the number of IKKγ molecules per cell was quantified in five tested cell lines to range from 4·10^5^ to 2·10^6^ molecules ^64^ which is only slightly higher than the inferred 9.8·10^4^ IKKγ molecules in our model. In conclusion, available *in vitro* studies support the predictions of our model regarding molecule numbers and proportions between ATM, PARP-1 and IKKγ as well as the input-output behaviour of ATM activation and NF-κB activation. In the future, it would be interesting to further analyse the impact of different molecule numbers and proportions and thereby assess differences in the response of IKK/ NF-κB to genotoxic stress at the level of cell types and individual cells.

Besides analysing the regulation of signalosome formation, which is the first signalling hub in the modelled network and was predicted to control the level of activated IKK complex in steady state, we also assessed the regulation of the second signalling hub. The parallel signalling merging in the second hub was predicted to control the initiation of activation of the IKK complex. Of note, the signal integration in the second hub forms a network motif termed coherent feedforward loop type 1, which has been shown to be able to delay the activation of the target component and filter transient inputs ^65^. By quantifying the impact of posttranslationally modified IKKγ (spIKKγ) and the ATM-TRAF6-TAK1 complex (ATT) on the readout IKK complex, we could demonstrate that ATT dominantly controls the onset of IKK complex activation (Fig 6). Thus, ATT determines the delay of IKK complex activation. ATM induces the formation of the TRAF6 complex and is also crucial for the activation of the transcription factor and tumour suppressor p53 which plays a fundamental role in the genotoxic stress response. As p53 induces the expression of Wip1, a phosphatase dephosphorylating and thereby inactivating ATM, the activity of p53 can modulate the activity of ATM ^66^ and therefore might be important at later time points in controlling the temporal response of NF-κB. For the second described characteristic of the feedforward motif, the filter for transient inputs, one can hypothesize that it prevents a premature activation of IKK/ NF-κB signalling upon minor DNA lesions. Poltz and Naumann ^67^ identified multiple coherent feedforward loops in a logical model of the DNA damage response. However, the majority of the identified feedforward loops were found for p53 signalling. The importance of this network motif for the regulation of p53 was demonstrated by Loewer et al., who showed that the filter prevents p53 activation in case of transient DNA damage ^68^.

Taken together, we here combined mechanistic insights and quantitative data of several experimental studies to develop a model for IKK/ NF-κB activation by genotoxic stress. The analysis of the model shows how the recognition of DNA double strand breaks by the two sensors PARP-1 and the MRN complex are integrated and transduced to the cytoplasm where the IKK complex and subsequently NF-κB is activated. Interestingly, our study reveals that the regulation properties of the pathway strongly depend on the irradiation dose. The pronounced impact of PARP-1 inhibition on IKK/ NF-κB activation indicates that the anti-apoptotic impact of NF-κB might affect the outcome of clinically applied PARP-1 inhibitors. This notion is in line with experimental observations reporting PARP-1 inhibition to cause a reduction in the viability of cells in an NF-κB-dependent manner ^26, 59, 69^. Therefore, a consideration of the activation of IKK/ NF-κB signalling by genotoxic stress could improve predicting the efficacy of PARP-1 inhibition and expand the application of inhibitors to tumour cells in which IKK/ NF-κB activity is upregulated or constitutively active. The identified coherent feedforward loop is another level of regulation besides signalosome formation and might be a possible point of cross-talk between p53 signalling and IKK/ NF-κB activity. Our model could be extended by interactions with the p53 signalling pathway and thereby serve as a starting point to gain a deeper understanding of cell fate decisions imposed by genotoxic stress. It will also be an important extension for disease models capturing the cellular response to oncogenic perturbations ^70, 71^.

## Methods

### Quantification of western blots and electrophoretic mobility shift assays

For the quantification of Western blots and electrophoretic mobility shift assay data, we used the software ImageJ ^72^. To determine the intensity of protein bands, the ImageJ in-built functions for polyacrylamide gel analysis was applied. The band intensity of the protein of interest was normalised to the corresponding protein band of the loading control for the Western blots.

### Model simulations and parameter inference

For simulations and parameter inference we used MATLAB (R2017b, The Mathworks Inc., Natick, MA) in combination with the open source toolbox Data2Dynamics ^73, 74^. For a detailed description, see Text S1, Section II.D.

### Sensitivity analysis

To identify processes with the strongest impact on inhibiting IKK/ NF-κB activity under genotoxic stress, we performed a sensitivity analysis for all kinetic model parameters by reducing each parameter individually by 90%. In order to investigate the sensitivity of ATM activation, we moreover reduced the two parameters k3 and k5 simultaneously by 90%. The activated IKK complex (pIKK) in stimulated steady state was chosen as readout for the analysis. The sensitivity coefficients (sc) for the specified parameters were calculated for various irradiation doses ranging from 1 Gy to 100 Gy by using the following formula:

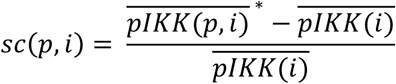

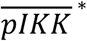 represents the level of activated IKK complex in the stimulated steady state for irradiation dose *i* upon the perturbation of a parameter *p*. 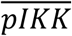 represents the level of activated IKK complex in the stimulated steady state without perturbation of parameter *p*. The most effective inhibition of IKK complex activation is characterised by parameter perturbations causing a reduction in the level 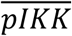 to a value of zero which results in a sensitivity coefficient of −1, given 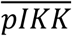 is nonzero. Parameter perturbations causing an increase in IKK complex activation can result in sensitivity coefficients higher than +1 as there is no upper limit for positive coefficients.

## Data and Code availability

The model will be publicly available in the BioModels data base ^75^ at https://www.ebi.ac.uk/biomodels/.

## Acknowledgments

F.K. was funded by a PhD fellowship of the graduate school ‘Computational Systems Biology’ of the German Research Foundation (DFG Graduiertenkolleg 1772). The project was supported by a grant from the German Federal Ministry of Education and Research BMBF (Project ProSiTu, 0316047A) to J.W. and C.S., and the e:Med-program of the German Ministry of Education and Research: SeneSys for iLymTx (grant number: 031L0189D) to J.W. The funders had no role in study design, data collection and analysis, decision to publish, or preparation of the manuscript.

## Author contributions

J.W. and C.S. designed the project. J.W., D.B and C.S. supervised the study. Data analysis, quantitative dynamic modelling, method development and model analysis were performed by F.K., Visualization was done by F.K. and J.W., M.W carried out the experiment. F.K., C.S. and J.W. wrote and all authors approved the manuscript.

