## Supplemental Text S1 for "A computational model of the DNA damage-induced IKK/ NF-κB pathway reveals a critical dependence on irradiation dose and PARP-1"

### I. Experimental data sets used for parameter inference and validation

For the calibration and validation of the developed model, we obtained available data sets covering various experimental assays and irradiation (IR) doses inducing DNA damage. Table S1 gives an overview of the data sets used in this study. We integrated data of different research groups and from different cell lines.

*Table S1: Overview of data sets used for calibration and validation of the model*

| Description | cell line | stimulus type | stimulus dose | reference |
|---|---|---|---|---|
| recruitment of PARP-1 | MEF | microirradiation | 405nm diode laser | [1] |
| ATM and MRN recruitment | U2OS, AT | γ-IR, X-ray | 0.8 – 68 Gy | [2,3] |
| ATM activation | HeLa, HepG2 | γ-IR | 40 Gy | [4,5] |
| signalosome formation | MEF | γ-IR | 80 Gy | [4] |
| TRAF6 complex formation | HeLa, HepG2 | γ-IR | 30, 40 Gy | [5] |
| validation (NF-κB) | HepG2 | γ-IR | 40 Gy | this study |

##### A. Modification of PARP-1

To quantify the recruitment dynamics of PARP-1 to DNA lesions, Mortusewicz et al.[1] measured the fluorescent intensity of GFP-labelled PARP-1 at the damage site upon inducing DNA damage using microirradiation in single cells. In the study, they measured the intensity of wild type (WT)-PARP-1 for two different time periods with a different temporal resolution (Fig S1, PARP(WT)). To investigate the effect of PARylated PARP-1 on the recruitment dynamics, they quantified the recruitment of an additional PARP-1 variant that is incapable of undergoing PARylation but still binds to the damage site (Fig S1, PARP(E988K)).

##### B. ATM and MRN recruitment

The recruitment dynamics of MRN and ATM to DNA lesions was investigated and quantified by Tobias et al.[3] and Kozlov et al.[2] using time-resolved single cell microscopy with GFP-labelled NBS1, a subunit of the MRN complex, and YFP-labelled ATM (Fig S1, MRN and ATM).

##### C. ATM activation

Hinz et al.[5] and Stilmann et al.[4] measured phosphorylated ATM in the nucleus and cytoplasm upon γ-IR using Western blots (Fig S1, pATM(n) and pATM(c)).

##### D. Signalosome formation

The formation and dissociation of the signalosome was analysed by Stilmann et al.[4] using co-immunoprecipitation assays and Western blotting of different components of the signalosome (Fig S1, sig).

##### E. TRAF6 complex formation

To capture signal propagation in the cytoplasm, we used Western blot data from Hinz et al.[5] tracking posttranslationally modified TRAF6, TAK1 as well as IKKγ at different time points upon γ-IR (Fig S1, pUbTRAF6, pTAK1 and mUbIKKγ).

##### F. Validation (NF-κB)

The activation of NF-κB was monitored using an electrophoretic mobility shift assay (Fig 2B).

#### II. Model description

##### A. Model structure

For the development of a quantitative model, we split the signalling pathway of genotoxic IKK/ NF-κB activation into smaller modules and created for each module multiple minimal models to investigate different mechanistic hypotheses[6]. Using mechanistic insights and experimental data from the studies presented in section I enabled us to identify the most appropriate model for each module. By merging the individual models, we created an overall model of genotoxic IKK/ NF-κB signalling that is presented in this study.

**Modelling DNA lesions.** To model the recruitment and binding of PARP-1 and MRN to DNA lesions, we implemented a double strand break by defining two binding sites for PARP-1 (BS_P) and two binding sites for MRN (BS_M). Introducing separate binding sites for PARP-1 and MRN prevents an exclusive binding of one of the two components to a binding site and is based on the study of Haince et al. in which PAR molecules, PARP-1 as well as MRE11, a subunit of the MRN complex, colocalize at a break site[7]. In the experiments that we used to calibrate and validate the model (see Table S1), DNA damage is induced by irradiation of cells for a defined time period. To represent a stimulus-dependent generation of binding sites, we defined the two parameters *kDNA_b* and *DNAb.* While *kDNA_b* determines the rate at which binding sites for PARP-1 and MRN are generated, *DNAb* controls the duration of the stimulus. The parameter *kDNA_b* is calculated by the following formula:

$$k_{DNAb} = r\ \frac{Gy}{s} \cdot 35 \frac{DSB}{Gy} \cdot 2 \frac{BS}{DSB} \qquad \text{(S0)}$$

In equation S0, *r* determines the rate of irradiation and is specific for an experiment. Based on the studies of Gulston et al., Loebrich et al. and Prise et al., it is assumed that 1 Gy irradiation causes the generation of 35 DNA double strand breaks[8–10]. Moreover, the term is multiplied by a factor of two, to account for the fact that one double strand break leads to two binding sites. For the experimental data capturing the modification of PARP-1 (Table S1, first line), the irradiation dose could not be determined. We therefore fitted the parameter *k_DNAb* for this particular data set (Table S2). For all other data sets, *k_DNAb* was calculated according to eq. S0. The parameter *DNAb* is set to one for a time period given by the corresponding experiment. After this period, *DNAb* is set to zero. The generation of binding sites is computed by multiplying *kDNA_b* with *DNAb* (see eq. S21 in Supplement, Part C Reaction rates).

**Modelling PARP-1 recruitment to DNA.** In the model (see Fig 1A for the reaction scheme, Supplement Part B and C for the ODE and reaction rates) we assume that PARP-1 is either directly recruited to the lesion by being attached to the binding site *BS_P* (PARPDSB_b, eq. S22) or indirectly (PARP_b, eq. S24) by the recruitment of already bound and PARylated PARP-1 proteins (parPARP1_b). Our hypothesis is derived from the elaborated study of Mortusewicz et al.[1] in which the recruitment dynamics of various PARP-1 variants was quantified. We used the recruitment dynamics of WT PARP-1 and a catalytically inactive PARP-1 variant (E988K) to test multiple hypotheses by fitting different variants of minimal models to the data[6]. The superior model variant was then selected for the overall model. To include the effect of the catalytically inactive PARP-1 variant in the model, we implemented parameter *AB* that is set to one for WT PARP-1 or zero for the inactive E988K variant (eq. S26 and S27).

**Modelling the recruitment of MRN and ATM to DNA lesions.** For the recruitment of the MRN complex to the binding site (BS_M) and the recruitment and activation of ATM, we included positive feedbacks (eq. S30 and S33) to capture the cyclic process of MRN and ATM recruitment to the damage site. Specifically, the phosphorylation of H2AX histones by recruited ATM proteins in proximity of the DNA lesion causes the recruitment of MDC1 and additional MRN complexes. In turn, further ATM molecules are recruited and activated. Thus, MRN and ATM maintain their own recruitment[11]. We implemented the positive feedbacks by Hill terms that allow reproducing the sigmoidal shaped time course of activated ATM upon DNA damage (Fig S1, pATM(n)). As ATM exists in cells as inactive homodimers or multimers that transform by auto-phosphorylation into active monomers[12], we set the Hill coefficient for both Hill terms to a value of two.

**Modelling the signalosome formation and modification of IKKγ.** Upon activation of ATM and PARylation of PARP-1 the signalosome is formed and results in the sumoylation and phosphorylation of IKKγ (eq. S35 and S36). While activated ATM (pATM) mediates the phosphorylation of IKKγ, the SUMO-1 ligase PIASy sumoylates IKKγ. As there was no experimental data available capturing the levels of PIASy over time, we did not include this component in the model.

**Modelling the cytoplasmic complex formation and activation of the IKK complex.** Based on the findings of Hinz et al., phosphorylated ATM as well as modified IKKγ translocate to the cytoplasm where ATM induces the K63-linked polyubiquitination of TRAF6 which in turn facilitates the recruitment and activation of kinase TAK1. The ubiquitin ligase cIAP1 is recruited as well and mediates the mono-ubiquitination of IKKγ. For simplicity and due to lack of sufficient data we neglected cIAP1 in the model. In the model, the translocation of phosphorylated ATM to the cytoplasm is lumped together with the binding and polyubiquitination of TRAF6 (see eq. S37). The binding and activation of TAK1 is modelled in eq. S38. Note that the processes describing the translocation of modified IKKγ into the cytoplasm, mono-ubiquitination, integration into IKK complexes and the activation of the IKK complex by TAK1-mediated phosphorylation of IKKβ are modelled as a single reaction (eq. S39).

**B. Ordinary differential equations**

Most variable names correspond to the labels in Fig. 1. For some variables abbreviated names are introduced:

| | |
|---|---|
| *AMRN_b* | ATM bound to the DNA bound MRN complex |
| *sig* | signalosome consisting of pATM, parPARP1 and IKKγ |
| *spIKKγ* | sumoylated and phosphorylated IKKγ |
| *AT* | complex consisting of pATM and TRAF6 |
| *ATT* | complex consisting of pATM and TRAF6 and pTAK1 |
| *pIKK* | activated IKK complex in which IKKβ is phosphorylated |

$$\frac{d(BS\_P(t))}{dt} = v1 - v2 + v3 \tag{S1}$$

$$\frac{d(PARP1(t))}{dt} = v3 + v5 + v9 + v16 - v2 - v4 \tag{S2}$$

$$\frac{d(PARP1DSB\_b(t))}{dt} = v2 - v3 - v6 \tag{S3}$$

$$\frac{d(PARP1\_b(t))}{dt} = v4 - v5 - v7 \tag{S4}$$

$$\frac{d(parPARP1\_b(t))}{dt} = v6 + v7 - v8 \tag{S5}$$

$$\frac{d(parPARP1(t))}{dt} = v8 - v9 - v15 \tag{S6}$$

$$\frac{d(BS\_M(t))}{dt} = v1 - v10 - v11 \tag{S7}$$

$$\frac{d(MRN(t))}{dt} = -v10 - v11 \tag{S8}$$

$$\frac{d(MRN\_b(t))}{dt} = v10 + v11 - v12 + v13 + v14 \tag{S9}$$

$$\frac{d(AMRN\_b(t))}{dt} = v12 - v13 - v14 \tag{S10}$$

$$\frac{d(ATM(t))}{dt} = -v12 \tag{S11}$$

$$\frac{d(pATM(t))}{dt} = v13 + v14 - v15 + v16 - v17 \tag{S12}$$

$$\frac{d(IKK\gamma(t))}{dt} = -v15 \tag{S13}$$

$$\frac{d(sig(t))}{dt} = v15 - v16 \quad \text{(S14)}$$

$$\frac{d(spIKK\gamma(t))}{dt} = v16 - v19 \quad \text{(S15)}$$

$$\frac{d(TRAF6(t))}{dt} = -v17 \quad \text{(S16)}$$

$$\frac{d(AT(t))}{dt} = v17 - v18 \quad \text{(S17)}$$

$$\frac{d(ATT(t))}{dt} = v18 \quad \text{(S18)}$$

$$\frac{d(TAK1(t))}{dt} = -v18 \quad \text{(S19)}$$

$$\frac{d(pIKK(t))}{dt} = v19 \quad \text{(S20)}$$

Conserved moieties:

| | | |
|---|---|---|
| PARP1_tot | = | *PARP*1 + *PARP*1*DSB_b* + *PARP*1*_b* + *parPARP*1*_b* + *parPARP*1 + *sig* |
| MRN_tot | = | *MRN* + *MRN_b* + *AMRN_b* |
| ATM_tot | = | *ATM* + *AMRN_b* + *pATM* + *sig* + *AT* + *ATT* |
| IKKγ_tot | = | *sig* + *spIKKγ* + *pIKK* |
| TRAF6_tot | = | *TRAF6* + *AT* + *ATT* |
| TAK1_tot | = | *TAK1* + *ATT* |

#### C. Reaction rates

$$v1 = k\_DNAb \cdot DNAb(t) \quad \text{(S21)}$$

$$v2 = kon \cdot PARP1(t) \cdot BS\_P(t) \quad \text{(S22)}$$

$$v3 = koff \cdot PARP1DSB\_b(t) \quad \text{(S23)}$$

$$v4 = kr \cdot PARP1(t) \cdot parPARP1\_b(t) \quad \text{(S24)}$$

$$v5 = koff2 \cdot PARP1\_b(t) \quad \text{(S25)}$$

$$v6 = vact \cdot PARP1DSB\_b(t) \cdot AB \quad \text{(S26)}$$

$$v7 = vact \cdot PARP1\_b(t) \cdot AB \quad \text{(S27)}$$

$$v8 = koff\_act \cdot parPARP1\_b(t) \quad \text{(S28)}$$

$$v9 = kdepar \cdot parPARP1(t) \quad \text{(S29)}$$

$$v10 = k1 \cdot MRN(t) \cdot BS\_M(t) \cdot \frac{MRN\_b^n(t)}{km + MRN\_b^n(t)} \quad \text{(S30)}$$

$$v11 = k4 \cdot MRN(t) \cdot BS\_M(t) \quad \text{(S31)}$$

$$v12 = k2 \cdot ATM(t) \cdot MRN\_b(t) \quad \text{(S32)}$$

$$v13 = k3 \cdot AMRN\_b(t) \cdot \frac{pATM^n(t)}{km2 + pATM^n(t)} \quad \text{(S33)}$$

$$v14 = k5 \cdot AMRN\_b(t) \quad \text{(S34)}$$

$$v15 = SFM\_k1 \cdot parPARP1(t) \cdot pATM(t) \cdot IKK\gamma(t) \quad \text{(S35)}$$

$$v16 = SFM\_k2 \cdot sig(t) \quad \text{(S36)}$$

$$v17 = TM\_k1 \cdot pATM(t) \cdot TRAF6(t) \quad \text{(S37)}$$

$$v18 = TM\_k2 \cdot AT(t) \cdot TAK1(t) \quad \text{(S38)}$$

$$v19 = TM\_k3 \cdot spIKK\gamma(t) \cdot ATT(t) \quad \text{(S39)}$$

#### D. Model parameters and fitting procedure

For parameter inference, we set the bounds of parameters to -10 and 3 on a $\log_{10}$ scale if not stated otherwise (Table S2). Variables and parameters in the model have units based on number of molecules (m) and seconds (s). For some of the parameters the boundaries were adapted to include information about experimentally quantified reaction rates from literature or to prevent numerical issues during simulation. For the half-life of PARylated PARP-1 (parPARP1) a time range of one to six minutes was published[13] which corresponds to an exponential decay rate of -1.9 and -2.6 $s^{-1}$ on a $\log_{10}$ scale. We therefore set the boundaries of the corresponding parameter *kdepar* to -4 and 0 $s^{-1}$ (Table S2). For the PARylation of PARP-1 a rate of 0.41 $s^{-1}$ and 5 $s^{-1}$ was reported[14,15]. The total concentrations of the conserved moieties ATM, IKK, MRN, TAK1 and TRAF6 were fitted and the boundaries were set to 2 and 7 on a $\log_{10}$ scale to ensure that the estimated values are in a physiological range. The total amount of PARP-1 was set to 187881 molecules in accordance to[16].

To infer parameter values, we used the MATLAB open source toolbox D2D[17,18]. The optimization is based on maximum likelihood estimation and the MATLAB in-built function lsqnonlin was used for minimizing the objective function. To prevent local minima, multi-start optimization was used. The parameter values of the best fit are given in table S2.

To assess the identifiability of fitted parameters, we used a profile likelihood-based approach[19] and applied it to each kinetic parameter of the model (Fig S3). The analysis shows that almost all estimated parameters are identifiable, i.e. there exists a finite range for the confidence interval of a parameter. In contrast, for the parameters MRN_tot, TAK1_tot, TRAF6_tot, kdepar and koff2 only the lower or upper limit of the confidence interval can be properly determined as the upper or lower boundary of the parameter value is reached. For example, for MRN_tot the lower limit of the confidence interval can be determined, but for the upper limit, the parameter boundary is reached at $10^7$ molecules. Overall, the results show that the data used for parameter estimation contains sufficient information for the given model to infer its parameter values.

**E. The total amount of modified IKKγ determines the level of activated IKK complex in steady state**

Based on the model structure and the implementation of processes, it is possible to comprehend that the amount of modified IKKγ (spIKKγ) produced over time corresponds to the amount of activated IKK complex in the stimulated steady state. This is due to i) the assumed irreversible translocation and integration of modified IKKγ into IKK complexes (parameter TM_k3 in Fig 1A and eq. S39 in II.C) and ii) the implementation of IKKγ as a conserved moiety and iii) a catalytic function of the cytoplasmic TRAF6 complex (ATT) for the activation of IKK (parameter TM_k3 in Fig 1A and eq. S39 in II.C), i.e. there is no mass flow from the TRAF6 complex to the IKK complex. As the cytoplasmic TRAF6 complex is not degraded and does not dissociate in the model, it cannot restrict the amount of modified IKKγ that is integrated into the IKK complex once the TRAF6 complex is formed.

**F. Defining activated IKK as readout for NF-κB activity**

For our analyses, we defined the activated IKK complex as a readout for NF-κB activity. The IKK complex is a direct activator of NF-κB as it phosphorylates and thereby initiates the degradation of IκBs which inhibit NF-κB. In addition, the IKK complex phosphorylates RelA/p65. Upon degradation of IκBs, NF-κB translocates into the nucleus and induces the expression of its target genes. Since we here aimed to model the initial activating processes that are specific for the genotoxic pathway we focused on the upstream processes leading to genotoxic stress-induced NF-κB activation. As NF-κB dynamics closely follow the dynamics of activated IKK (Fig 2B) we selected the activated IKK complex as the readout of our model.

*Table S2: Parameter description with estimated parameter values*

| label | $log_{10}$ value | $log_{10}$ values of boundaries | description |
|---|---|---|---|
| SFM_k1 | -7.0 $m^{-2}\cdot s^{-1}$ | [-10, -2] | formation of signalosome |
| SFM_k2 | -1.3 $s^{-1}$ | [-10, 3] | dissociation of signalosome |
| TM_k1 | -4.5 $m^{-1}\cdot s^{-1}$ | [-10, 3] | poly-ubiquitination of TRAF6 and binding of pATM |
| TM_k2 | -10.2 $m^{-1}\cdot s^{-1}$ | [-12, 3] | activation of TAK1 and recruitment to TRAF6 complex |
| TM_k3 | -3.8 $m^{-1}\cdot s^{-1}$ | [-10, 3] | integration of IKKγ into IKK complex, mono-ubiquitination of IKKγ and phosphorylation of IKKβ |
| ATM_tot | 3.9 m | [ 2, 7] | total concentration of ATM |
| IKKγ_tot | 5.0 m | [ 2, 7] | total concentration of IKKγ |
| MRN_tot | 6.9 m | [ 2, 7] | total concentration of MRN |
| PARP1_tot | 5.3 m | [ -, -] | total concentration of PARP-1 |
| TAK1_tot | 6.2 m | [ 2, 7] | total concentration of TAK1 |
| TRAF6_tot | 2.0 m | [ 2, 7] | total concentration of TRAF6 |
| k1 | -9.0 $m^{-1}\cdot s^{-1}$ | [-10, 3] | positive feedback of chromatin-bound MRN on MRN recruitment |
| k2 | -5.2 $m^{-1}\cdot s^{-1}$ | [-10, 3] | recruitment of ATM to MRN |
| k3 | -3.2 $s^{-1}$ | [-10, 1] | positive feedback of pATM on activation of ATM |
| k4 | -9.5 $m^{-1}\cdot s^{-1}$ | [-10, 3] | recruitment of MRN to DNA lesion |
| k5 | -3.4 $s^{-1}$ | [-10, 1] | activation of ATM and its dissociation from MRN |
| k_DNAb | 3.6 $m\cdot s^{-1}$ | [ 2, 4] | generation of binding sites |
| kdepar | 0.0 $s^{-1}$ | [ -4, 0] | dePARylation of PARP-1 |
| km | 3.8 $m^{2}$ | [ -4, 9] | parameter of Hill term for positive feedback of MRN |
| km2 | 1.9 $m^{2}$ | [ -4, 9] | parameter of Hill term for positive feedback of ATM |
| koff | -1.4 $s^{-1}$ | [-10, 3] | dissociation of PARP-1 from DNA lesion |
| koff2 | 2.9 $s^{-1}$ | [-10, 3] | dissociation of PARP-1 from periphery of lesion |
| koff_act | 1.1 $s^{-1}$ | [-10, 3] | dissociation of PARylated PARP-1 from chromatin |
| kon | -8.9 $m^{-1}\cdot s^{-1}$ | [ -9, -2] | binding of PARP-1 to lesion |

| | | | |
|---|---|---|---|
| kr | -2.1 $m^{-1} \cdot s^{-1}$ | [ -9, 3] | PARylated PARP-1-induced recruitment of PARP-1 to periphery of lesions |
| n | $log_{10}(2)$ | [ -, -] | Hill coefficient for positive feedback of MRN and ATM |
| vact | 0.8 $s^{-1}$ | [-0.4, 1] | automodification of PARP-1 |

Abbreviations: m: molecules, s: seconds
