## Supplemental Figures for "A computational model of the DNA damage-induced IKK/ NF-κB pathway reveals a critical dependence on irradiation dose and PARP-1"

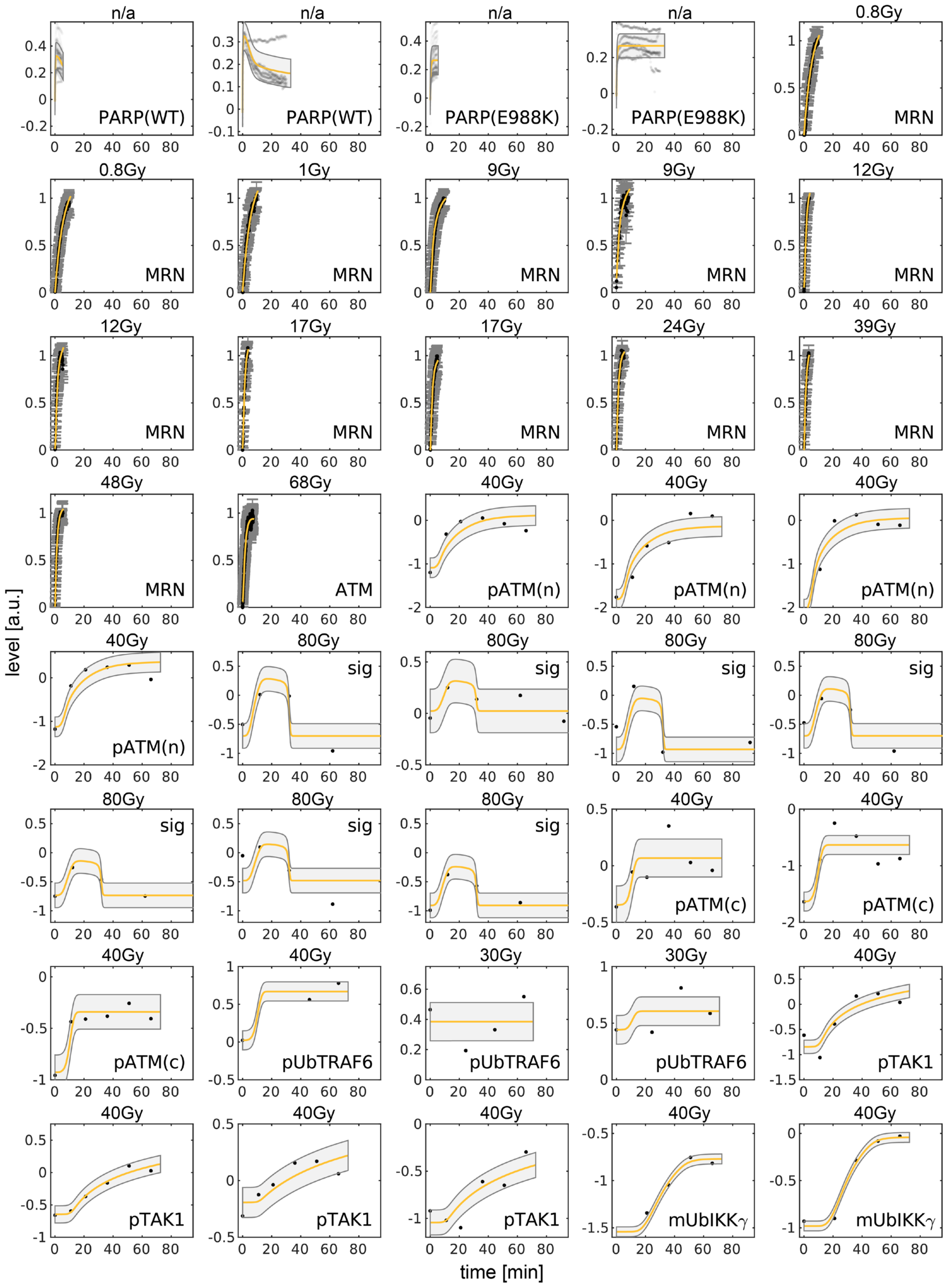


**Fig S1. Experimental data and simulations of calibrated model.**

Dots represent experimental data and the yellow line represents simulation data. The grey area surrounding the yellow line shows the fitted standard deviation. Otherwise the standard deviation is represented by grey bars which is true for the MRN and ATM recruitment data. For the recruitment data of the PARP-1 WT and mutant two similar data sets exist which differ in the temporal resolution and time period. For those experiments, the irradiation dose is not available (n/a). For 0.8, 12, and 17 Gy for the recruitment data of the MRN complex, additional biological replicates exist. Details about the experimental data and settings are given in Text S1, section I. The estimated parameter values for the best fit are shown in Text S1 (table S2). Abbreviation: sig: signalosome.

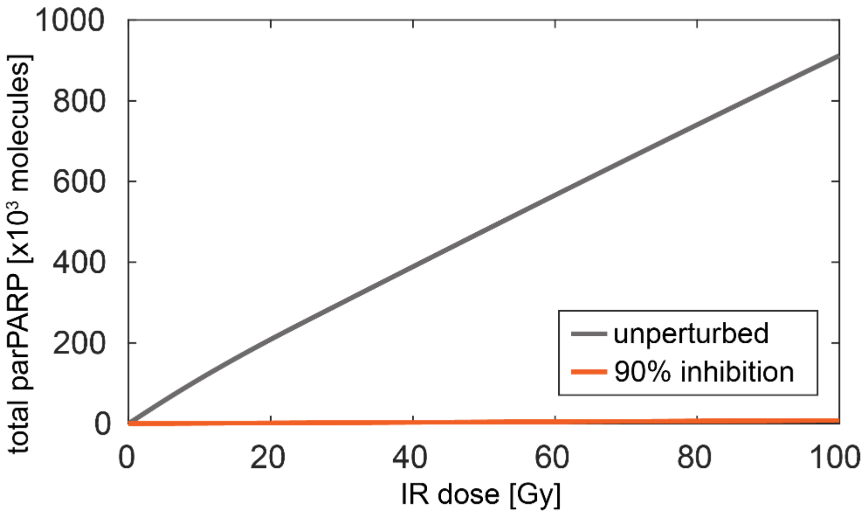


**Fig S2. Influence of the irradiation dose and inhibition on total amount of PARylated PARP-1.**

The total amount of PARylated PARP-1 was quantified by integrating the influx of PARylated PARP-1 (koff_act) for various irradiation doses with and without 90% inhibition of PARylation (vact).

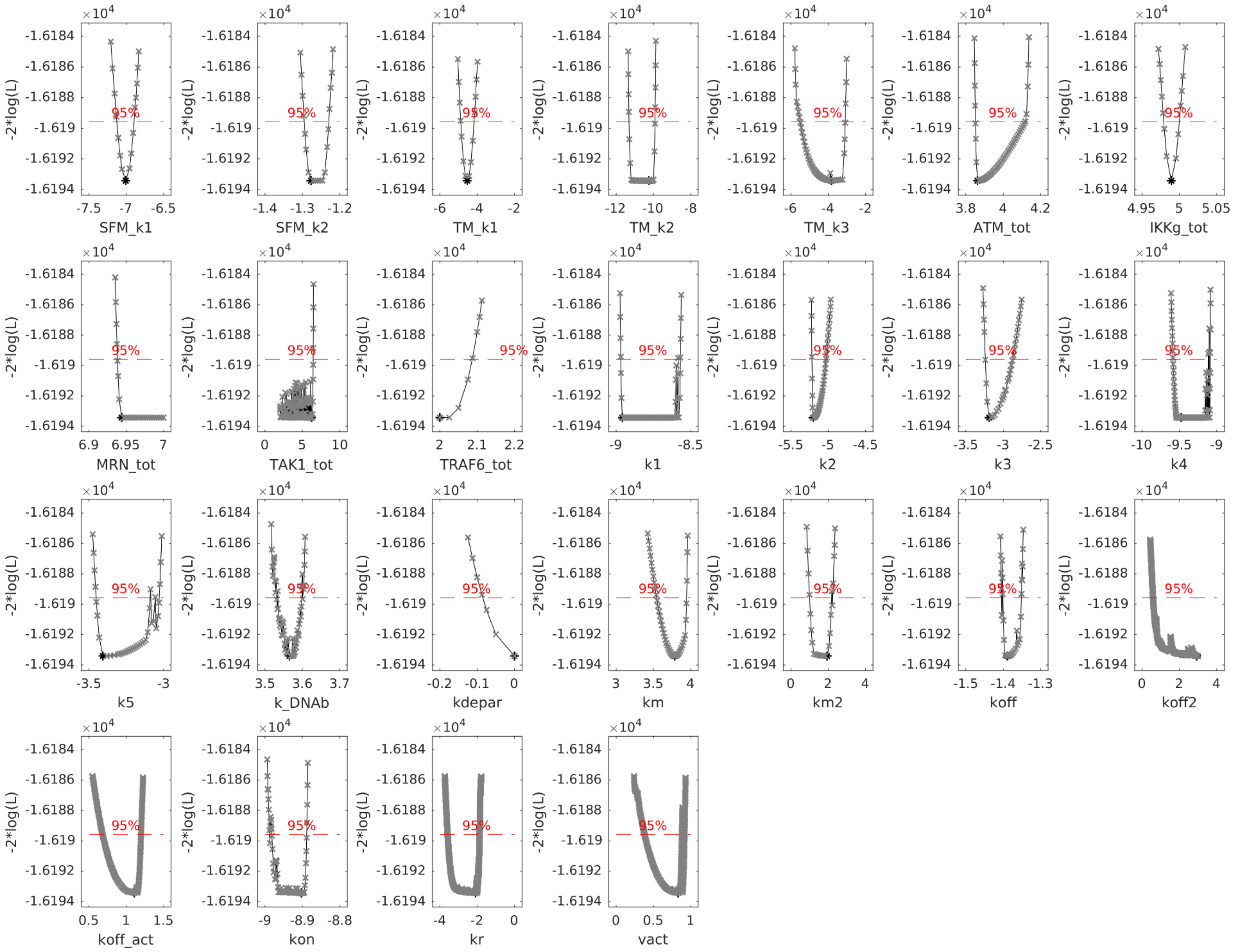

×10^4
-2*log(L)
-1.6184
-1.6186
-1.6188
-1.619
-1.6192
-1.6194
95%
SFM_k1
SFM_k2
TM_k1
TM_k2
TM_k3
ATM_tot
IKKg_tot
MRN_tot
TAK1_tot
TRAF6_tot
k1
k2
k3
k4
k5
k_DNAb
kdepar
km
km2
koff
koff2
koff_act
kon
kr
vact

**Fig S3. Profile likelihoods for estimated model parameters.**

The black star represents the estimated parameter value of the best fit and the corresponding fit quality given as negative log likelihood. To compute the profile likelihood, the specified parameter is fixed to an increased or decreased value compared to the calibrated value and the remaining parameters are refitted. The resulting log likelihood is represented by a grey x. To identify the 95% confidence interval of a parameter, a threshold for the log likelihood is calculated based on a $\chi^2$ distribution with one degree of freedom (dashed red line).
